# Gsx2 is involved in specification of neurons in the inferior olivary nuclei from Ptf1a-expressing neuronal progenitors in zebrafish

**DOI:** 10.1101/2020.03.16.993402

**Authors:** Tsubasa Itoh, Miki Takeuchi, Marina Sakagami, Kazuhide Asakawa, Koichi Kawakami, Takashi Shimizu, Masahiko Hibi

## Abstract

Neurons in the inferior olivary nuclei (IO neurons) send climbing fibers to Purkinje cells to elicit functions of the cerebellum. IO neurons and Purkinje cells are derived from neural progenitors expressing the proneural gene *ptf1a*. In this study, we found that the homeobox gene *gsx2* was co-expressed with *ptf1a* in IO progenitors in zebrafish. Both *gsx2* and *ptf1a* zebrafish mutants showed a strong reduction or loss of IO neurons. The expression of *ptf1a* was not affected in *gsx2* mutants and *vice versa*. In IO progenitors, the *ptf1a* mutation increased apoptosis whereas the *gsx2* mutation did not, suggesting that *ptf1a* and *gsx2* are independently regulated and have distinct roles. The fibroblast growth factors (Fgf) 3/8a and retinoic acid signals negatively and positively, respectively, regulated *gsx2* expression and thereby the development of IO neurons. *mafba* and *hox* genes are at least partly involved in the Fgf- and retinoic acid-dependent regulation of IO neuronal development. Our results indicate that *gsx2* mediates the rostro-caudal positional signals to specify the identity of IO neurons from *ptf1a*-expressing neural progenitors.

**Summary:** The homeobox gene *gsx2* mediates rostro-caudal positional signaling to specify the identify of neurons in the inferior olivary nuclei from neural progenitors expressing the proneural gene *ptf1a*.

## INTRODUCTION

In an initial step of the formation of neural circuits, neurons are differentiated from their neural/neuronal progenitors. The fate of neurons is thought to be determined by inductive signals that their progenitors receive depending on the position of the neural/neuronal progenitors. By receiving positional signals, neural progenitors express transcription factors, such as basic helix-loop-helix (bHLH) and homeobox transcription factors, that function in the differentiation and specification of neurons. Elucidation of these transcription factors and their regulation is a prerequisite to understanding the mechanism of neural circuit formation. Cerebellar neural circuits provide a good model for revealing positional signal-dependent neurogenesis.

In both mammals and teleosts, such as zebrafish, cerebellar neural circuits are composed of several types of neurons, including granule cells (GCs) and Purkinje cells (PCs) in the cerebellum, as well as neurons in the inferior olivary nuclei (IO neurons) in the caudal hindbrain. PCs receive axons from GCs (parallel fibers) and IO neurons (climbing fibers: CFs), then integrate the input information to elicit the cerebellum’s functions. PCs then send output to the outside of the cerebellum via efferent neurons, which are eurydendroid cells in zebrafish. In zebrafish, a portion of GCs also project their axons to dendrites of crest cells, which are PC-like cells located in the medial octavolateral nucleus (MON) of the dorsal hindbrain (Bae et al., 2009; Hashimoto and Hibi, 2012; Hibi and Shimizu, 2012). These neurons are derived from the hindbrain, which contains seven compartments collectively referred to as the rhombomere (r). PCs and IO neurons are generated from neural progenitors in the dorsal ventricular zone of the hindbrain. Analyses of mutants and cell lineage tracing for the bHLH-type proneural gene pancreas transcription factor 1a (*Ptf1a*) in mice revealed that PCs and IO neurons are derived from *Ptf1a*-expressing progenitors and *Ptf1a* is required for generation of these neurons in mice (Hoshino et al., 2005; Yamada et al., 2007). Lineage tracing with transgenic (Tg) zebrafish also showed that PCs and IO neurons are derived from *ptf1a*-expressing (*ptf1a*^+^) progenitors in zebrafish (Kani et al., 2010). Inhibitory neurons in the dorsal cochlear nuclei (DCN neurons) were also shown to be derived from *Ptf1a^+^* neural progenitor cells in r2-5 in mice (Fujiyama et al., 2009). These reports indicate that despite being present in the dorsal ventricular zone of all rhombomeres, *ptf1a*^+^ progenitors generate different types of neurons depending on their position. At r1, PCs are generated from *ptf1a*^+^ progenitors and migrate dorsally whereas in the caudal hindbrain, IO neurons are generated from *ptf1a*^+^ progenitors and migrate ventrally (reviewed in (Hashimoto and Hibi, 2012; Hoshino, 2012)). However, it is not clear yet how positional signals are interpreted to specify neurons from *ptf1a^+^* progenitors and what molecules control differentiation and/or specification of IO neurons.

Rostro-caudal patterning of the caudal hindbrain is controlled by gradients of fibroblast growth factor 3/8a (Fgf3/8a) and retinoic acid (RA) signals (Dupe and Lumsden, 2001; Gavalas and Krumlauf, 2000; Marin and Charnay, 2000; Maves et al., 2002; Maves and Kimmel, 2005). During early neurogenesis in zebrafish, *fgf3/8a* are expressed at r4 and are required for fate determination of r5 and r6 (Maves et al., 2002; Shimizu et al., 2006; Walshe et al., 2002; Wiellette and Sive, 2004). In the aldehyde dehydrogenase 1 family, the A2 gene (*aldh1a2*) encodes an enzyme which is required for the production of RA and is expressed in forming somites during early neurogenesis (Begemann et al., 2001; Grandel et al., 2002). RA signaling is required for the formation of r5-7 and the anterior spinal cord (Begemann et al., 2001; Grandel et al., 2002; Niederreither et al., 2000; Shimizu et al., 2006). These data suggest that Fgf and RA signal gradients in which Fgf and RA signals are high at r4 and the caudal-most hindbrain, respectively, play important roles in specifying neurons in the caudal hindbrain. Although Fgf and RA signals regulate the expression of *mafba*, *knox20*, and *hox* cluster genes (Ghosh et al., 2018; Marin and Charnay, 2000; Walshe et al., 2002), it is largely unknown which genes function downstream of these signals to regulate neuronal specification.

In mice, the homeobox gene *Gsx2* (formerly *Gsh2*) is expressed in the ventral telencephalon and is involved in fate determination of lateral ganglionic eminence (LGE) progenitor cells, which give rise to striatal neurons and olfactory bulb interneurons (Corbin et al., 2000; Hsieh-Li et al., 1995; Szucsik et al., 1997; Waclaw et al., 2009; Yun et al., 2003). *Gsx2*, which is also expressed in the ventricular zone of the spinal cord and hindbrain, is involved in the generation of a subset of hindbrain neurons and dorsal interneurons in the spinal cord of mice (Kriks et al., 2005; Mizuguchi et al., 2006; Satou et al., 2013). However, the role of *gsx2* in the development of IO neurons has not been elucidated. In this report, we show that *gsx2* is co-expressed with *ptf1a* in neuronal progenitors of IO neurons and is required for the development of IO neurons in zebrafish. *gsx2* expression is regulated through transcriptional networks that are activated by Fgf and RA signals in the caudal hindbrain. Our results reveal that *gsx2* mediates the rostro-caudal positional signals to control the identity of IO neurons.

## RESULTS

### *gsx2* is co-expressed in IO progenitors that express *ptf1a*

To reveal the mechanisms that generate IO neurons, we sought genes expressed in IO progenitors in the dorso-caudal hindbrain. A previous study with a BAC transgenic fish *TgBAC(gsx2:LOXP-Tomato-LOXP-GFP)* (hereafter referred to as *Tg(gsx2:RFP)*) suggested that *gsx2* is expressed in the caudal hindbrain (Satou et al., 2013). Thus, we focused on *gsx2* and *ptf1a*, which is also involved in the generation of IO neurons in mice (Yamada et al., 2007). We found that *gsx2* was expressed in the dorsal part of the caudal hindbrain and the ventral telencephalon in an early stage (2, 3 days post fertilization: dpf) of zebrafish larvae (Fig. 1A, B, E, F). As reported previously, *ptf1a* was expressed in the dorsal ventricular zone of all rhombomeres (Fig. 1C, D, G, H). To examine *gsx2*-expressing cells in detail, we compared RFP expression in *Tg(gsx2:RFP)* larvae (gsx2:RFP) with GFP expression in gSAIzGFFM35A;*Tg(5xUAS:EGFP)* (Fig. 2I-L) and *Tg(ptf1a:EGFP)* larvae (ptf1a:GFP, Fig. 1M-T). gSAIzGFFM35A is a Gal4 trap line that harbors a Gal4 gene in the exon of *mafba* (previously known as *valentino* or *kreisler*) and can drive UAS-mediated GFP expression at r5 and 6 (referred to as mafba^GFF^;UAS:GFP, Fig. 1I-L) (Asakawa and Kawakami, 2018). The gsx2:RFP^+^ and mafba^GFF^;UAS:GFP^+^ cells did not overlap and were separated adjacently in the caudal hindbrain at an early larval stage (3 dpf, Fig. 1I-L), indicating that *gsx2* is specifically expressed in r7 of the hindbrain and the rostral spinal cord. As reported previously (Kani et al., 2010), ptf1a:GFP expression was detected in neural progenitors located in the dorsal ventricular zone of all rhombomeres at an early larval stage (5 dpf, Fig. 1M, N, Q). gsx2:RFP expression was detected in the dorsal ventricular zone of only the caudal hindbrain (Fig. 1M, O, Q). gsx2:RFP and ptf1a:GFP were co-expressed in cells in the dorsal ventricular zone (Fig. 1M, Q). Since GFP and RFP (Tomato) are relatively stable, they can be used for lineage tracing in zebrafish (Kani et al., 2010). Although *gsx2* and *ptf1a* were not detected in the ventral hindbrain (Fig. 1E-H), gsx2:RFP and ptf1a:GFP were detected in IO neurons in the ventral hindbrain in 5-dpf larvae (Fig. 1R, S, T). These results indicate that *gsx2* is co-expressed with *ptf1a* in IO progenitors.

**Fig. 1.**
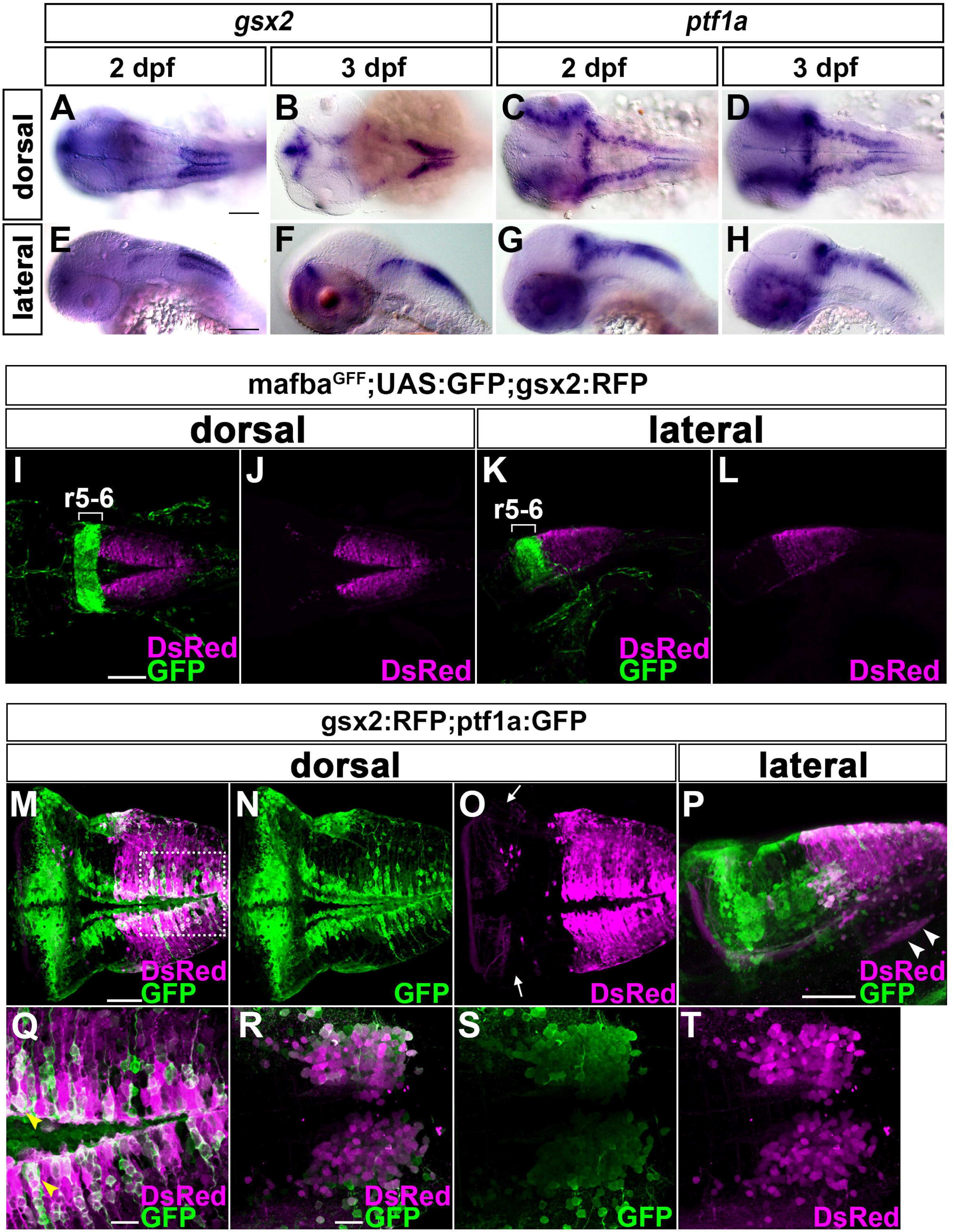
*gsx2* and *ptf1a* are co-expressed in IO progenitors. (A-B, E-F) Expression of *gsx2* in 2- and 3-dpf larvae. (C-D, G-H) Expression of *ptf1a* in 2- and 3-dpf larvae. (I-L) 3-dpf mafba^GFF^;UAS:GFP;gsx2:RFP larvae (*n*=2) were immunostained with anti-DsRed (RFP, magenta) and anti-GFP (green) antibodies. r5-6: rhombomeres 5 and 6. (M-T) 5-dpf gsx2:RFP;ptf1a:GFP larvae were immunostained with anti-DsRed (magenta) and anti-GFP (green) antibodies. (Q) Highly magnified view of the dotted box in M (dorsal optical section). (R-T) Expression of GFP and/or RFP in the IO region. Arrows and arrowhead indicate CFs and IO neurons, respectively. Yellow arrowheads indicate both GFP^+^ and RFP^+^ cells. Scale bars: 100 µm in A (applies to A-H) and I (applies to I to L); 50 µm in M (applies to M-O) and P; 20 µm in Q and R (applies to R-T).

**Fig. 2.**
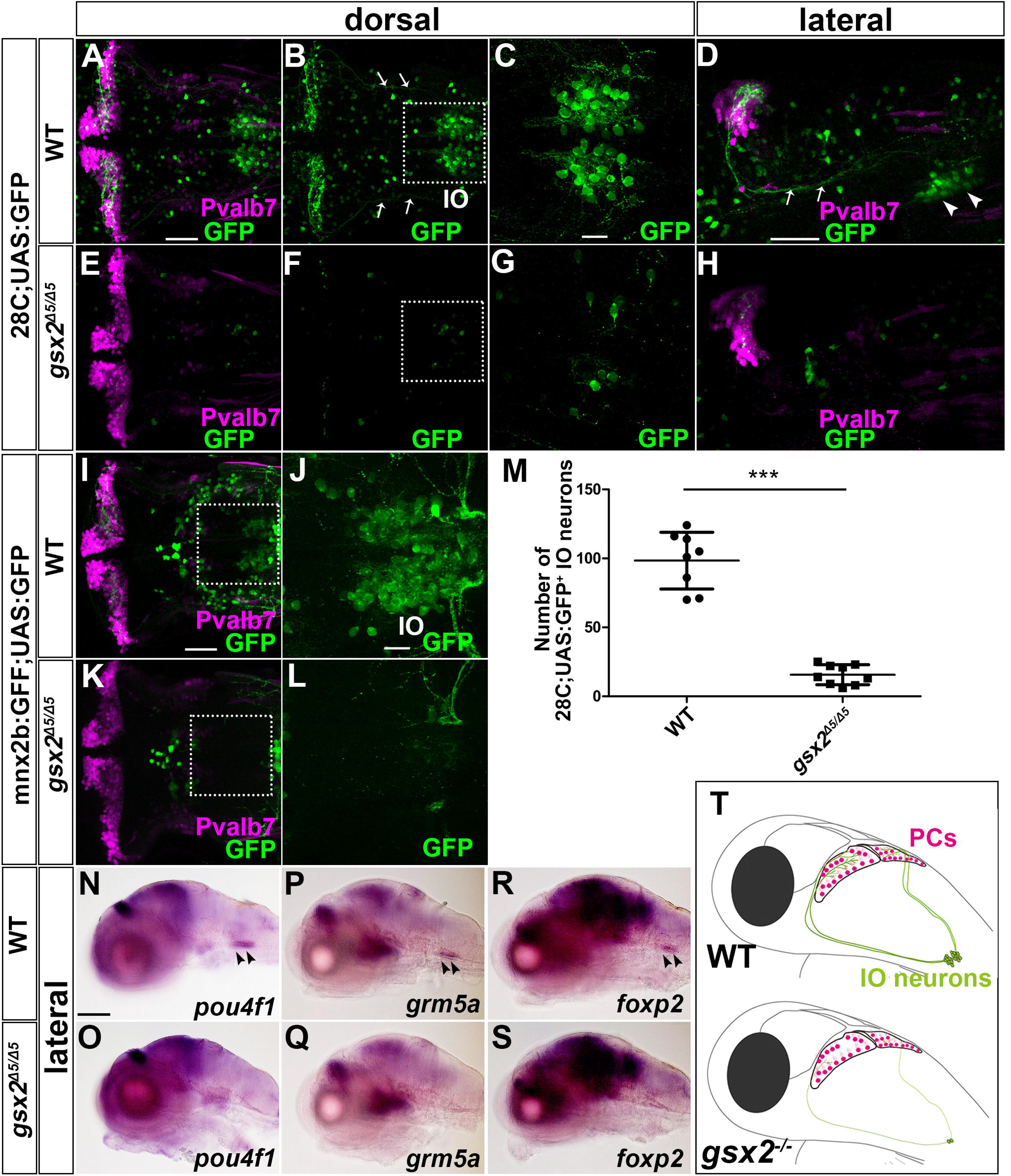
Strong reduction or loss of IO neurons in *gsx2* mutants. (A-H) 5-dpf wild-type (WT, A-D, *n*=10) and *gsx2^Δ5/Δ5^* (E-H, *n*=10) hspGFFDMC28C;UAS:GFP (28C;UAS:GFP) larvae which were immunostained with anti-Pvalb7 (magenta) and anti-GFP (green) antibodies. Arrows and arrowhead indicate CFs and IO neurons, respectively. (C, G) Highly magnified views of the boxes in B and F. (I-L) 5-dpf WT (I-J *n*=6) and *gsx2 ^Δ5/Δ5^* (K-L *n*=7) mnx2b:GFF;UAS:GFP larvae. (J, L) Highly magnified views of the boxes in I and K. (M) Number of 28C;UAS:GFP^+^ IO neurons in WT and *gsx2^Δ5/Δ5^* larvae. 28C;UAS:GFP^+^ IO neurons were significantly reduced in *gsx2* mutant larvae compared to control larvae. \*\*\**p* < 0.001 (Student’s *t*-test). (N-S) Expression of endogenous IO neuronal markers in WT and *gsx2^Δ5/Δ5^*. (N-O) Expression of *pou4f1* in WT (*n*=4) and *gsx2^Δ5/Δ5^* (*n*=4). (P-Q) Expression of *grm5a* in WT (*n*=3) and *gsx2^Δ5/Δ5^* (*n*=2). (R-S) Expression of *foxp2* in WT (*n*=5) and *gsx2^Δ5/Δ5^* (*n*=5). (T) Illustration of PCs and IO neurons in WT and *gsx2* mutant larvae. Scale bars: 100 µm in N (applies to N-S); 50 µm in A (applies to A-B, E-F), D (applies to H), and I (applies to K); 20 µm in C (applies to G) and J (applies to L).

### The *gsx2* mutation leads to a strong reduction or loss of IO neurons

To reveal the roles of Gsx2 in IO neuronal development, we generated *gsx2* mutants by the CRISPR/Cas9 method. The *gsx2* mutants harbor a 5-bp (*gsx2*^*Δ5*^) or 8-bp (*gsx2^Δ8^*) deletion in exon 1 of the *gsx2* gene that introduces a premature stop (Fig. S1). Although *gsx2* mRNA was not reduced in the *gsx2* mutant larvae (Fig. S2), the Gsx2 proteins of the putative mutants lack the homeodomain (Fig. S1) and the mutations are likely null alleles. To analyze the development of IO neurons in *gsx2* mutants, we used the Tg line hspGFFDMC28C;*Tg(5xUAS:EGFP)* (28C;UAS:GFP) which expresses GFP in IO neurons (Takeuchi et al., 2015). GFP^+^ somata were located in the ventral hindbrain and they extended their axons (CFs) to PCs, which were marked with anti-parvalbumin7 (Pvalb7) antibody in control 5-dpf Tg larvae (Fig. 2A-D, Fig. S3). In contrast, GFP^+^ somata were rarely detected and GFP^+^ CFs were not observed in *gsx2* mutants (Fig. 2E-H, M, Fig. S3). In addition to 28C;UAS:GFP, we found that the *TgBAC(mnx2b:GFF);Tg(5xUAS:EGFP)* line also expressed GFP in a portion of IO neurons in addition to motoneurons (mnx2b:GFF;UAS:GFP) (Asakawa et al., 2013) (Fig. 2I, J). Similar to 28C;UAS:GFP^+^ cells, a strong reduction or loss of mnx2b:GFF;UAS:GFP^+^ IO neurons was observed in 5-dpf *gsx2^Δ5^* mutant larvae (Fig. 2K, L). We further examined endogenous markers of IO neurons. *pou4f1* (also known as *brn3a*) and a glutamate receptor, metabotropic 5a (*grm5a*), are expressed in zebrafish IO neurons (Bae et al., 2009; Haug et al., 2013). *foxp2* is expressed in IO neurons of mice (Ferland et al., 2003; Fujita and Sugihara, 2012). Whereas the expression of these markers was detected in IO neurons of wild-type (WT) larvae, it was not observed in *gsx2* mutant larvae (Fig. 2N-S, Fig. S3), indicating a strong reduction or absence of IO neurons in *gsx2* mutants. These data reveal that Gsx2 is required for IO neuronal development (Fig. 2T).

### The *ptf1a* mutation leads to a strong reduction or loss of IO neurons

We next analyzed the roles of *ptf1a* in the development of IO neurons by generating *ptf1a* mutants by the CRISPR/Cas9 method. The *ptf1a* mutants have a 4-bp (*ptf1a^Δ4^*) deletion or 11-bp insertion (*ptf1a^+11^*) in exon 1 (Fig. S1). Although the *ptf1a* mutant larvae retained *ptf1a* mRNA (Fig. S2), the putative mutant Ptf1a proteins lack the bHLH domain and the mutations were likely null alleles. The *ptf1a* mutant larvae displayed a complete loss of IO neurons that express GFP in 28C;UAS:GFP or mnx2b:GFF;UAS:GFP Tg lines (Fig. 3E-H, K, L, Fig. S3). Furthermore, the IO markers *pou4f1*, *grm5a*, and *foxp2* were not detected in *ptf1a* mutant larvae (Fig. 3M-R, Fig. S3). These results are consistent with the defective development of IO neurons in *Ptf1a* mutant mice (Yamada et al., 2007). In addition to IO neurons, the *ptf1a* mutants showed a reduction – but not loss – of PCs and crest cells, which were positive for Pvalb7, as was shown previously (Bae et al., 2009) (Fig. 3E, S4). These data suggest that even though PCs, crest cells, and IO neurons are derived from *ptf1a*-expressing progenitors, *ptf1a* functions redundantly with other genes during the development of PCs and crest cells while *ptf1a* is essential for the development of IO neurons in zebrafish (Fig. 3S).

**Fig. 3.**
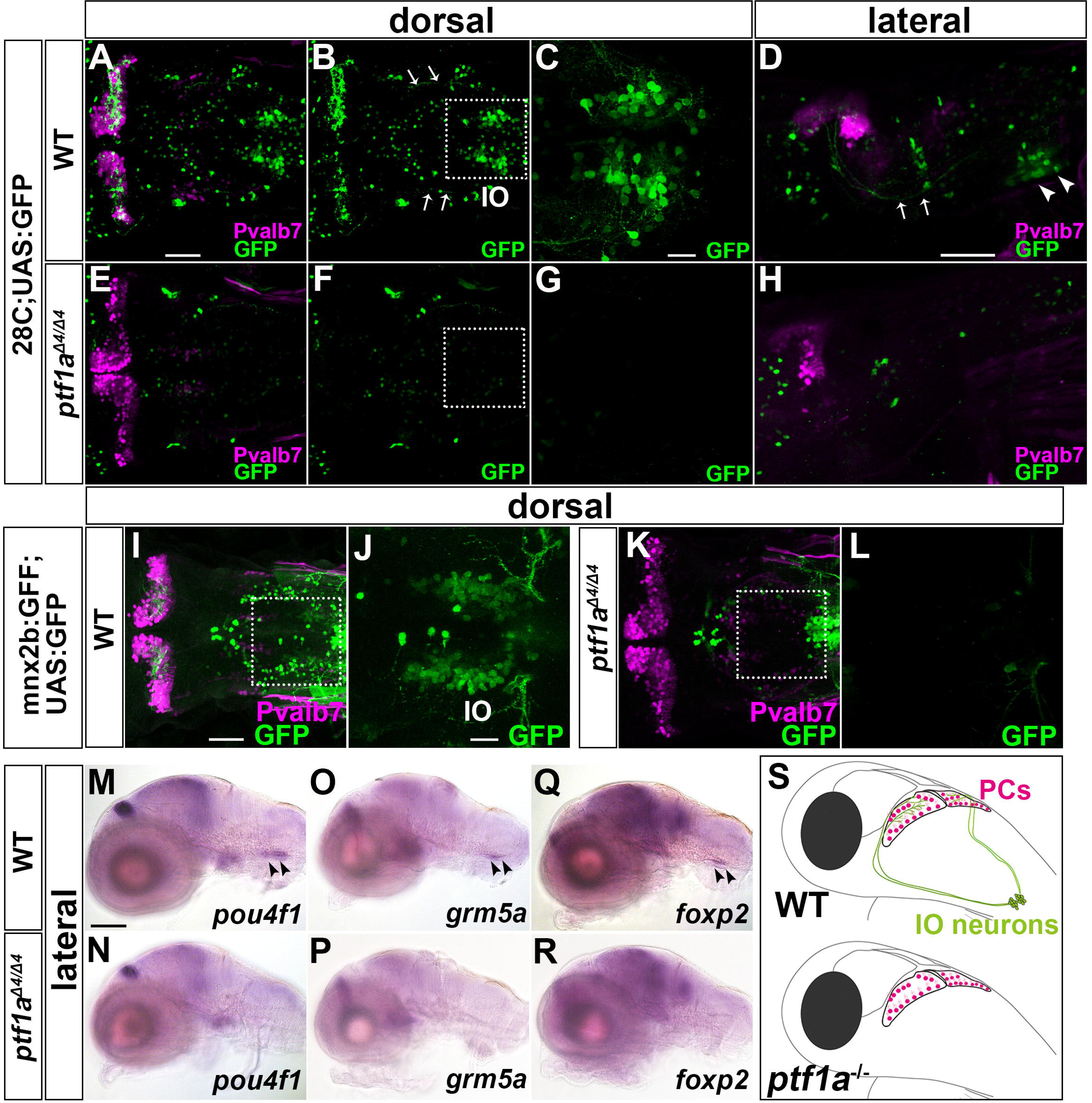
Loss of IO neurons in *ptf1a* mutants. (A-H) 5-dpf WT (A-D, *n*=5) and *ptf1a^Δ4/Δ4^* (E-H, *n*=5) 28C;UAS:GFP larvae were immunostained with anti-Pvalb7 (magenta) and anti-GFP (green) antibodies. Arrows and arrowhead indicate CFs and IO neurons, respectively. (C, G) Highly magnified views of the boxes in B and F. (I-L) 5-dpf WT (I-J *n*=2) and *ptf1a^Δ4/Δ4^* (K-L *n*=4) mnx2b:GFF;UAS:GFP larvae. (J, L) Highly magnified views of the boxes in I and K. (M-N) Expression of *brn3a* in 5-dpf WT (*n*=4) and *ptf1a^Δ4/Δ4^* (*n*=4) larvae. (O-P) Expression of *grm5a* in 5-dpf WT (*n*=4) and *ptf1a^Δ4/Δ4^* (*n*=3) larvae. (Q-R) Expression of *foxp2* in 5-dpf WT (*n*=5) and *ptf1a^Δ4/Δ4^* (*n*=5) larvae. (S) Illustration of Purkinje cells and IO neurons in WT and *ptf1a* mutant larvae. Scale bars: 100 µm in M (applies to M-R); 50 µm in A (applies to A-B, E-F), D (applies to H) and I (applies to K); 20 µm in C (applies to G) and J (applies to L).

### Independent regulation and distinct role of *gsx2* and *ptf1a* in IO progenitors

We examined the relationship between *gsx2* and *ptf1a* in IO neuronal development. The expression of *ptf1a* and *gsx2* was not affected in *gsx2* and *ptf1a* mutant larvae, respectively, during the development of IO neurons (Fig. 4). These results indicate that *ptf1a* and *gsx2* are independently regulated in the caudal hindbrain. We further analyzed apoptosis by immunohistochemistry with an anti-cleaved caspase3 antibody since *Ptf1a* mouse mutants show an increase in apoptotic cells in the caudal hindbrain (Yamada et al., 2007). As in *Ptf1a* mutant mice, the *ptf1a* zebrafish mutant larvae showed an increase in the number of cleaved-caspase3^+^ cells (Fig. 5A-H, Q). In contrast, there was no significant increase in cleaved-caspase3^+^ cells in *gsx2* mutant larvae (Fig. 5I-P, R). Furthermore, gsx2:RFP expression was retained in cells migrating from the dorsal progenitor domain in *gsx2* mutant larvae (Fig. S5), suggesting that *gsx2*^+^ progenitors remained undifferentiated or differentiated into other neurons in *gsx2* mutants. Our findings indicate that *ptf1a* and *gsx2* play different roles in IO neuronal development.

**Fig. 4.**
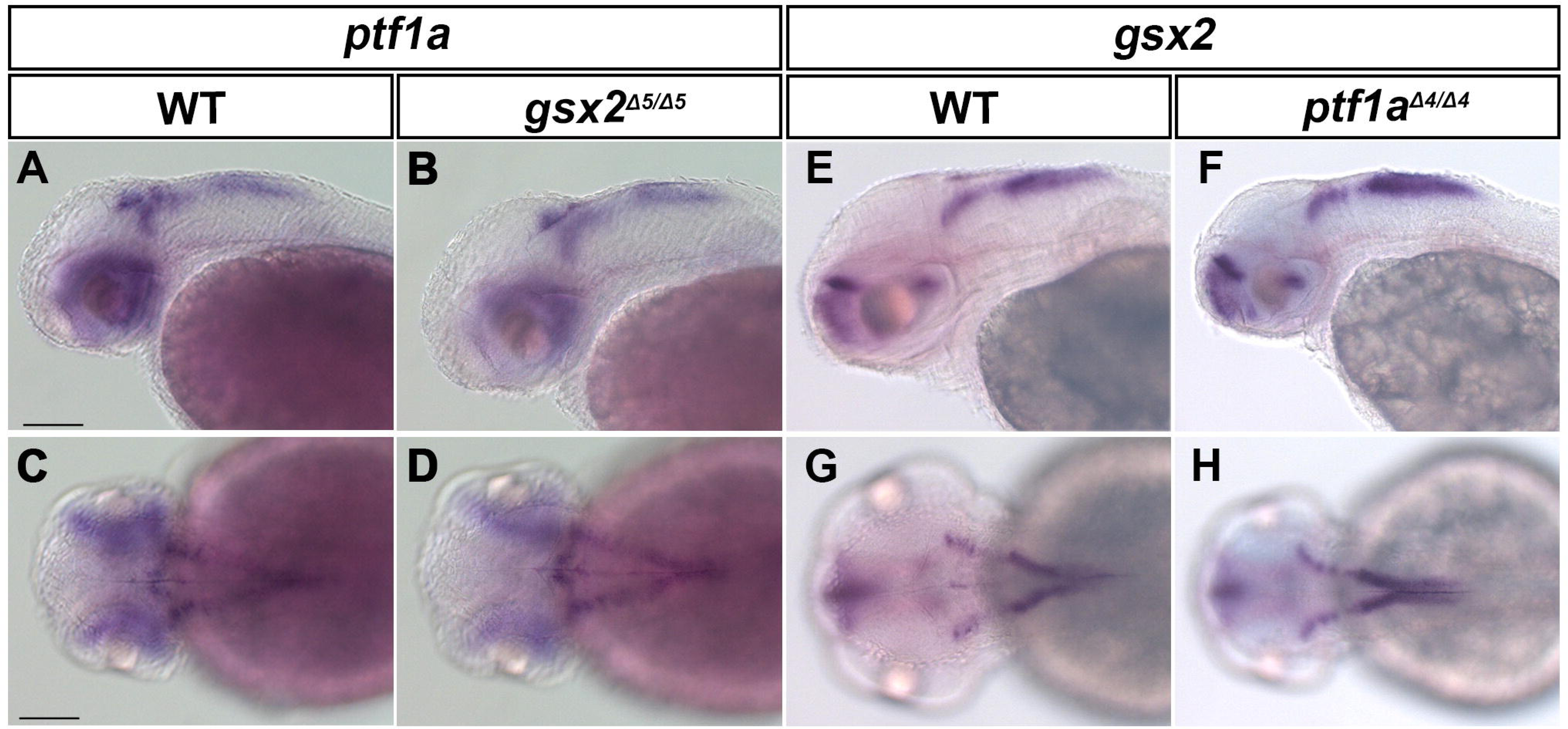
Expression of *ptf1a* and *gsx2* was not affected in *gsx2* and *ptf1a* mutants. (A-D) Expression of *ptf1a* in 2-dpf WT (A, C, *n*=2) and *gsx2* mutant (B, D, *n*=4) larvae. (E-H) Expression of *gsx2* in 2-dpf WT (E, G, *n*=2) and *ptf1a* mutant (F, H, *n*=3) larvae. Scale bars: 100 µm in A (applies to A-B, E-F) and C (applies to C-D, G-H).

**Fig. 5.**
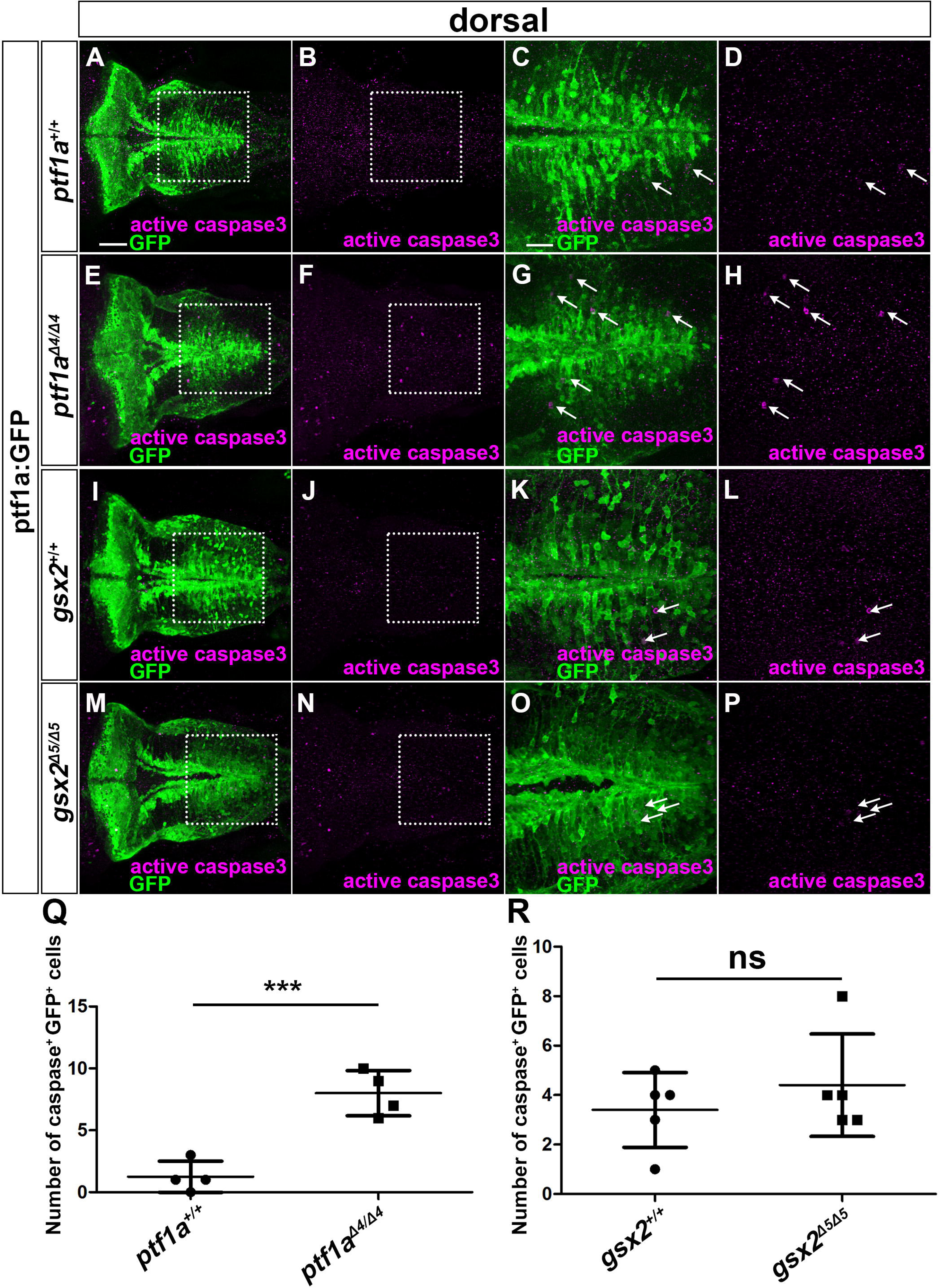
Apoptosis occurs in *ptf1a* but not *gsx2* mutants. (A-P) 5-dpf *ptf1a*^+/+^ (A-D, *n*=4), *ptf1a^Δ4/Δ4^* (E-H, *n*=4), *gsx2^+/+^* (I-L, *n*=5), and *gsx2^Δ5/Δ5^* (M-P, *n*=5) ptf1a:GFP larvae were stained with anti-GFP (green) and anti-cleaved caspase3 (magenta) antibodies. (C, G, K, O) Highly magnified views of the boxes in A, E, I and M. (D, H, L, P) Highly magnified views of the boxes in B, F, J and N. Arrows indicate both GFP^+^ and cleaved caspase3^+^ cells. (Q-R) Number of GFP^+^ and cleaved caspase3^+^ cells in the caudal hindbrain in WT, *ptf1a*, and *gsx2* mutant. ns: not significant; \*\*\**p* < 0.001 (Student’s *t*-test for Q and R). Scale bars: 50 µm in A (applies to A-B, E-F, I-J, M-N); 20 µm in C (applies to C-D, G-H, K-L, O-P).

### *gsx2* expression is negatively regulated by Fgf signaling and Mafba

Gradients of Fgf and RA signals are involved in the formation and patterning of the caudal hindbrain (Dupe and Lumsden, 2001; Gavalas and Krumlauf, 2000; Marin and Charnay, 2000; Maves et al., 2002). We inhibited Fgf and RA signals in order to investigate the regulation of *gsx2* expression. Antisense morpholino (MO)-mediated inhibition of *fgf3* did not significantly increase gsx2:RFP expression or the number of 28C;UAS:GFP^+^ IO cells compared to WT (Fig. 6C, D, M, R-T, AA), whereas *fgf8a* mutant larvae showed a slight increase in gsx2:RFP expression and the number of 28C;UAS:GFP^+^ IO neurons (Fig. 6E, F, M, U-W, AA). Inhibition of *fgf8a* and *fgf3* (injection of *fgf3* MO into *fgf8a* mutants: *fgf8a^ti282a/ti282a^*;*fgf3*^MO^) further increased gsx2:RFP expression and the number of 28C;UAS:GFP^+^ IO neurons (Fig. 6G, H, M, X-Z). Consistent with this finding, inhibition of the Fgf signal by SU5402, which is a chemical inhibitor of Fgf receptor tyrosine kinases (Mohammadi et al., 1997), led to an expansion of gsx2:RFP expression (Fig. 6K, L, N). Our findings suggest that the Fgf signal suppressed *gsx2* expression and thereby suppressed IO neuronal fate.

**Fig. 6.**
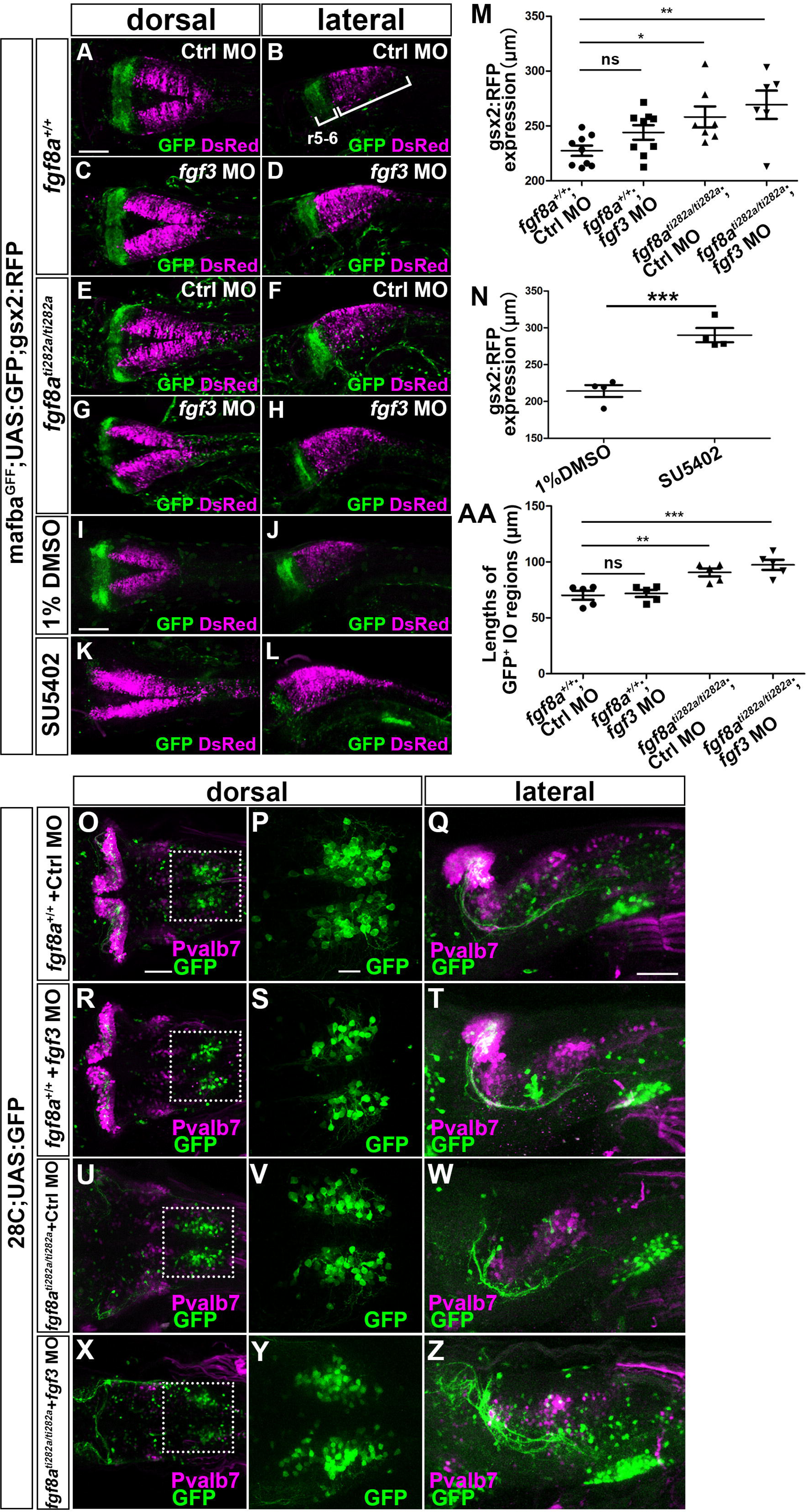
Fgf signal suppresses *gsx2* expression and IO neuronal development. (A-H) 3-dpf control (*fgf8a*^+/+^) or *fgf8a^ti282a/ti282a^* mafba^GFF^;UAS:GFP;gsx2:RFP larvae that received an injection of control MO or *fgf3* MO (*fgf3*^MO^) were stained with anti-RFP (magenta) and anti-GFP (green) antibodies. (I-L) mafba^GFF^;UAS:GFP;gsx2:RFP were treated with 1% DMSO (I, J, *n*=4) or 200 µM SU5402 (K, L, *n*=4) from 6 to 22 hpf, fixed at 3 dpf, and stained with anti-RFP (magenta) and anti-GFP (green) antibodies. (M, N) Length of gsx2:RFP^+^ hindbrain region (*gsx2* expression) in *fgf8a*^+/+^ (*n*=9), *fgf8a*^+/+^;*fgf3*^MO^ (*n*=9), *fgf8a ^ti282a/ti282a^* (*n*=7), and *fgf8a ^ti282a/ti282a^*; *fgf3*^MO^ (*n*=6) larvae (M), and in DMSO- (*n*=4) or SU5402-treated (*n*=4) larvae (N). (O-Z) Expression of GFP (green) and Pvalb7 (magenta) in control (*fgf8*^+/+^) or *fgf8a ^ti282a/ti282a^* 28C;UAS:GFP larvae received an injection of control MO or *fgf3* MO (*fgf3*^MO^). (P, S, V, and Y) Highly magnified views of the boxes in O, R, U, and X. (AA) Length of the 28C;UAS:GFP^+^ IO region in *fgf8a*^+/+^ (*n*=5), *fgf8a*^+/+^;*fgf3*^MO^ (*n*=5), *fgf8a ^ti282a/ti282a^* (*n*=5), and *fgf8a ^ti282a/ti282a^*; *fgf3*^MO^ (*n*=5) larvae. (AA) Length of GFP^+^ IO regions. r5-6: rhombomere 5 and 6. ns: not significant; \**p* < 0.05; \*\**p* < 0.01; \*\*\**p* < 0.001 (Student’s *t*-test for N and one-way ANOVA with Dunnett’s Multiple Comparison Test for M and AA). Scale bars: 100 µm in A (applies to A-H) and I (applies to I-L); 50 µm in O (applies to O, R, U, and X) and Q (applies to Q, T, W, and Z); 20 µm in P (applies to P, S, V, and Y).

mafba^GFF^;UAS:GFP expression was reduced when the Fgf signal was inhibited (Fig. 6G, H, K, L). We further analyzed the roles of *mafba* in *gsx2* expression and in the development of IO neurons (Fig. 7). In *mafba^GFF/GFF^* mutant larvae (Asakawa and Kawakami, 2018), rostral gsx2:RFP expression expanded slightly compared to control heterozygotes, and gsx2:RFP^+^ cells were observed in the mafba^GFF^;UASGFP^+^ region, corresponding to r5/6 (Fig. 7D-F). Furthermore, 28C;UAS:GFP^+^ IO neurons increased significantly in *mafba^b337/b337^* mutant larvae (Fig. 7J-L, N, O). These data suggest that Mafba is at least partly involved in the Fgf signal-mediated suppression of *gsx2* expression and IO neuronal development.

**Fig. 7.**
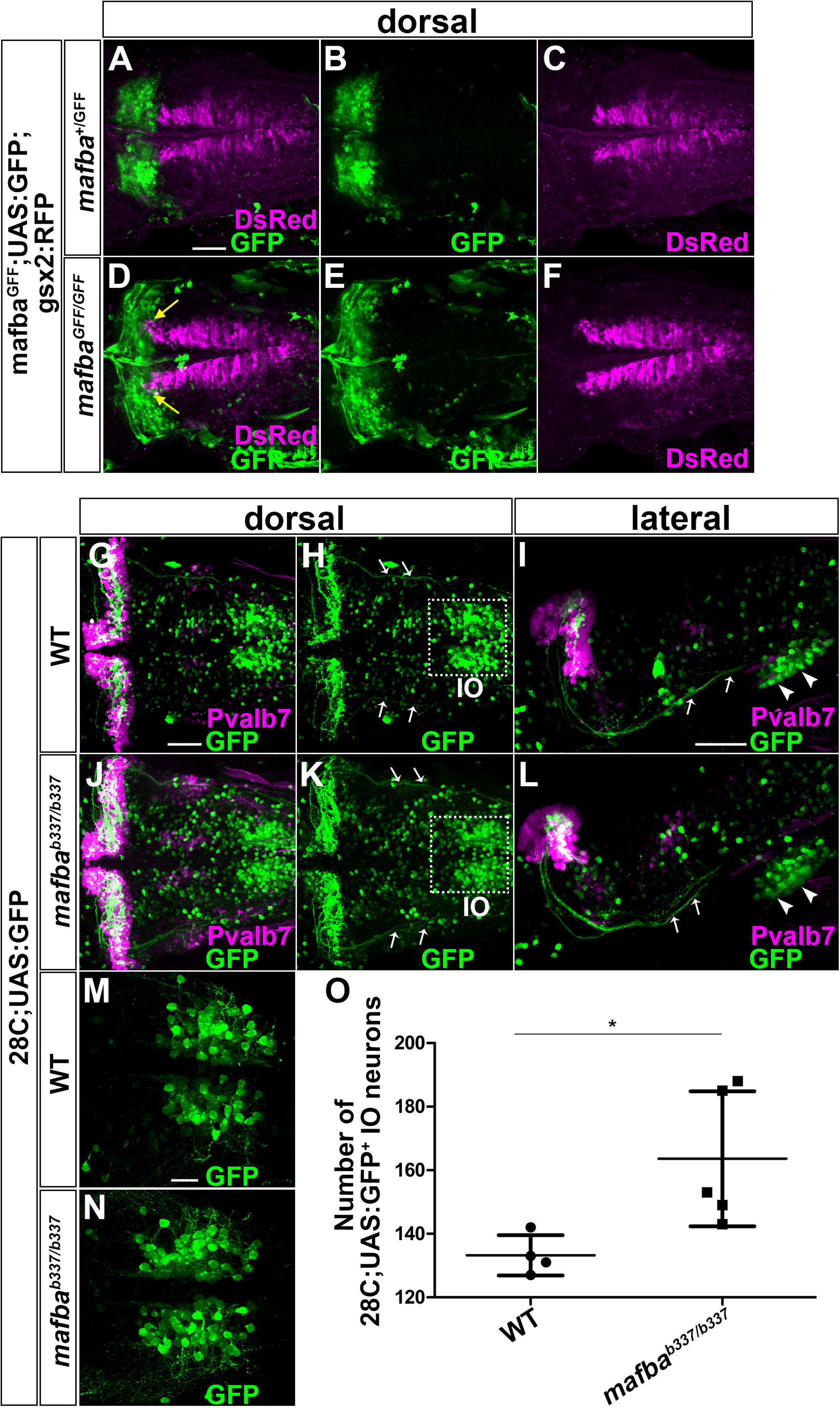
Mafba negatively controls *gsx2* expression and IO neuronal development. (A-F) 3-dpf *mafba^+/GFF^* (A-C, *n*=2) and *mafba*^GFF/GFF^ (D-F, *n*=5);UAS:GFP;gsx2:RFP larvae were stained with anti-RFP (magenta) and anti-GFP (green) antibodies. Yellow arrows indicate that *gsx2* expression is extend to rhombomeres 5 and 6. (G-N) 5-dpf WT and *mafba^b337/b337^* 28C;UAS:GFP larvae were stained with anti-GFP (green) and anti-Pvalb7 (magenta) antibodies. Arrows and arrowhead indicate CFs and IO neurons, respectively. (O) Number of GFP^+^ IO neurons in WT and *mafba^b337/b337^*. \**p* < 0.05 (Student’s *t*-test). Scale bars: 50 µm in A (applies A to F) and G (applies to G, H, J, and K) and I (applies to I and L); 20 µm in M (applies to M and N).

### *gsx2* expression is positively regulated by the RA-*hox* cascade

We inhibited the RA signal by a MO against *aldh1a2*, which recapitulates the phenotypes of an *aldh1a2* mutant (Begemann et al., 2001), or an Aldh1a2 inhibitor diethylaminobezaldehyde (DEAB) (Russo, 1997). Inhibition of the RA signal with both MO and DEAB resulted in reduced gsx2:RFP expression but higher mafba^GFF^;UASGFP expression (r5/6, Fig. 8A-J). It also reduced the number of 28C;UAS:GFP^+^ IO neurons (Fig. 8L, N, P, R, W, X) and expression of the IO neuronal marker *pou4f1* (Fig. 8S-V). The reduction was stronger in DEAB-treated larvae than in *aldh1a2* morphants, consistent with a previous report that showed that DEAB-treated larvae showed more severe phenotypes than *aldh1a2* mutants (Begemann et al., 2004). Our data indicate that the RA signal positively controls *gsx2* expression in the caudal hindbrain, and is also involved in IO neuronal development.

**Fig. 8.**
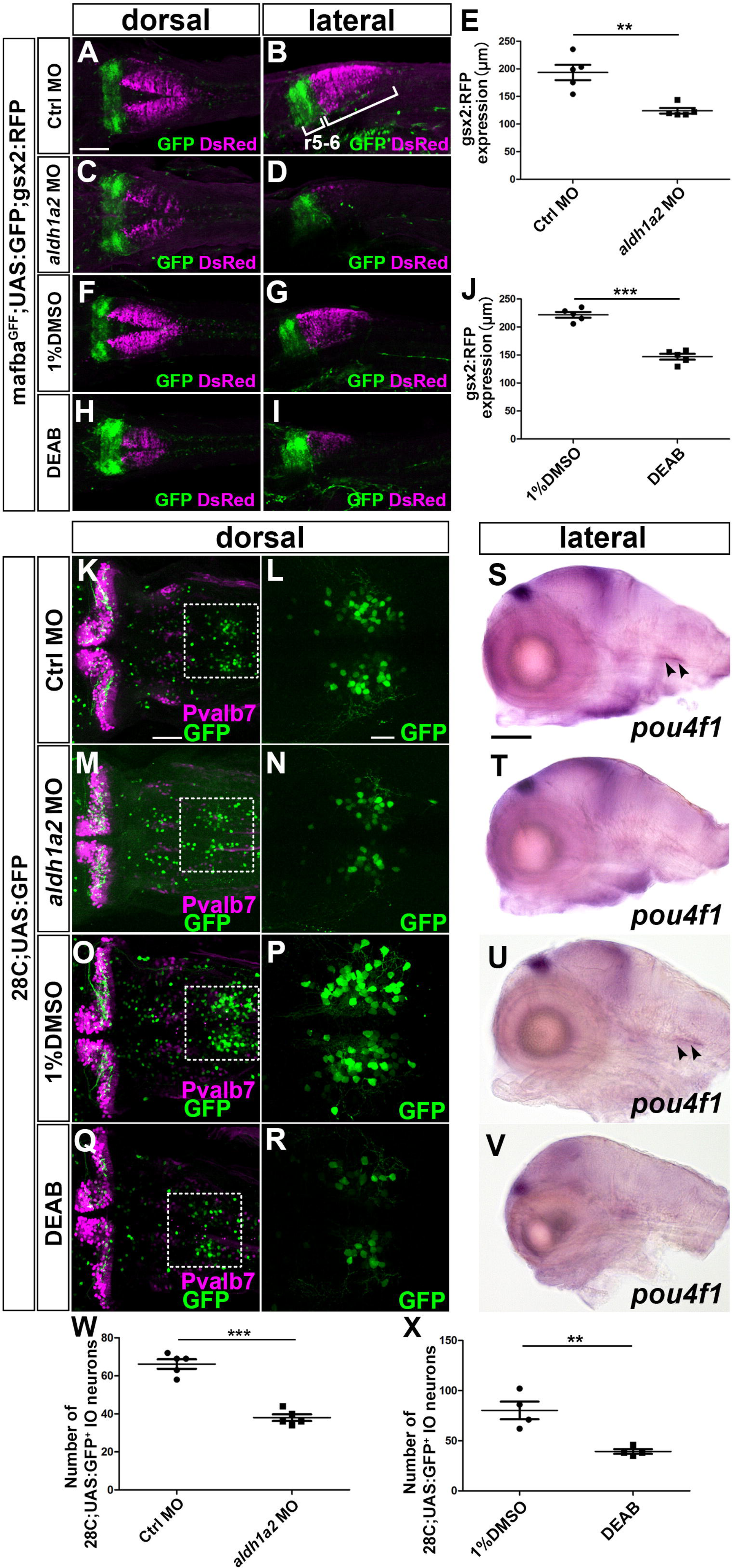
RA signal positively controls *gsx2* expression and IO neuronal development. (A-D) 3-dpf control morphant (A-B, *n*=5) and *aldh1a2* morphant (C-D, *n*=5) mafba^GFF^;UAS;GFP;gsx2:RFP larvae were stained with anti-RFP (magenta) and anti-GFP (green) antibodies. (E) Length of gsx2:RFP expression in the hindbrain in control and *aldh1a2* morphants. (F-I) mafba^GFF^;UAS;GFP;gsx2:RFP larvae were treated with 1% DMSO (F-G, *n*=5) and 0.25µM DEAB (H-I, *n*=5) from 6 to 22 hpf, fixed at 3 dpf, and stained with anti-GFP and anti-RFP antibodies. (J) Length of gsx2:RFP expression in the hindbrain of larvae treated with DMSO or DEAB. (K-R) 5-dpf control morphant (K, L, *n*=5) and *aldh1a2* morphant (M, N, *n*=5), DMSO (O, P, *n*=5) or DEAB-treated (Q, R, *n*=5) 28C;UAS:GFP larvae were stained with anti-Pvalb7 (magenta) and anti-GFP (green) antibodies. (L, N, P, R) Highly magnified views of the boxes in K, M, O, and Q. (S-V) Expression of *pou4f1* in 5-dpf control morphant (*n*=5), *aldh1a2* morphant (*n*=5), DMSO- (*n*=3) or DEAB-treated (*n*=3). (W) Number of 28C;UAS:GFP^+^ IO neurons in 5-dpf control morphant and *aldh1a2* morphant larvae. (X) Number of 28C;UAS:GFP^+^ IO neurons in 5-dpf larvae treated with DMSO and DEAB. \*\**p* < 0.01; \*\*\**p* < 0.001 (Student’s *t*-test). Arrows and arrowhead indicate CFs and inferior olivary nuclei, respectively. Scale bars: 100 µm in A (applies to A-D and F-I) and S (applies to S-V); 50 µm in K (applies to K, M, O, and Q); 20 µm in L (applies to L, N, P, and R).

The RA signal controls the expression of *hox* genes in the caudal hindbrain (Ghosh et al., 2018; Marin and Charnay, 2000). To address the roles of *hox* genes in IO neuronal development, we examined *gsx2* expression and IO neurons in larvae defective of Pbx2 and Pbx4, which are co-factors in the rostral Hox protein (Waskiewicz et al., 2002). The *pbx2/pbx4* morphant larvae recapitulate the phenotypes of maternal zygotic *pbx2* mutants in which *pbx4* inhibition was mediated by MO, and in which r2-6 acquired the r1 state (Maves et al., 2009). In *pbx2/4* morphant larvae, ectopic PCs were observed caudally but crest cells were present normally (Fig. 9G). In contrast, gsx2:RFP expression, the number of 28C;UAS:GFP^+^ IO neurons, and *pou4f1* expression were strongly reduced or absent in *pbx2/4*-deficent larvae (Fig. 9C, D, G, H, J, K). To address the roles of *hox* genes, we further ectopically expressed *hoxb4a* since it is expressed in r7 and in the spinal cord. Injection of *hoxb4a* mRNA expanded gsx2:RFP expression in the caudal hindbrain (Fig. 9N, O). Although *hoxb4a* might not be the only *hox* gene regulating caudal-most hindbrain identify, our data suggest that the RA-*hox* cascade plays a major role in controlling *gsx2* expression and subsequent IO neuronal development.

**Fig. 9.**
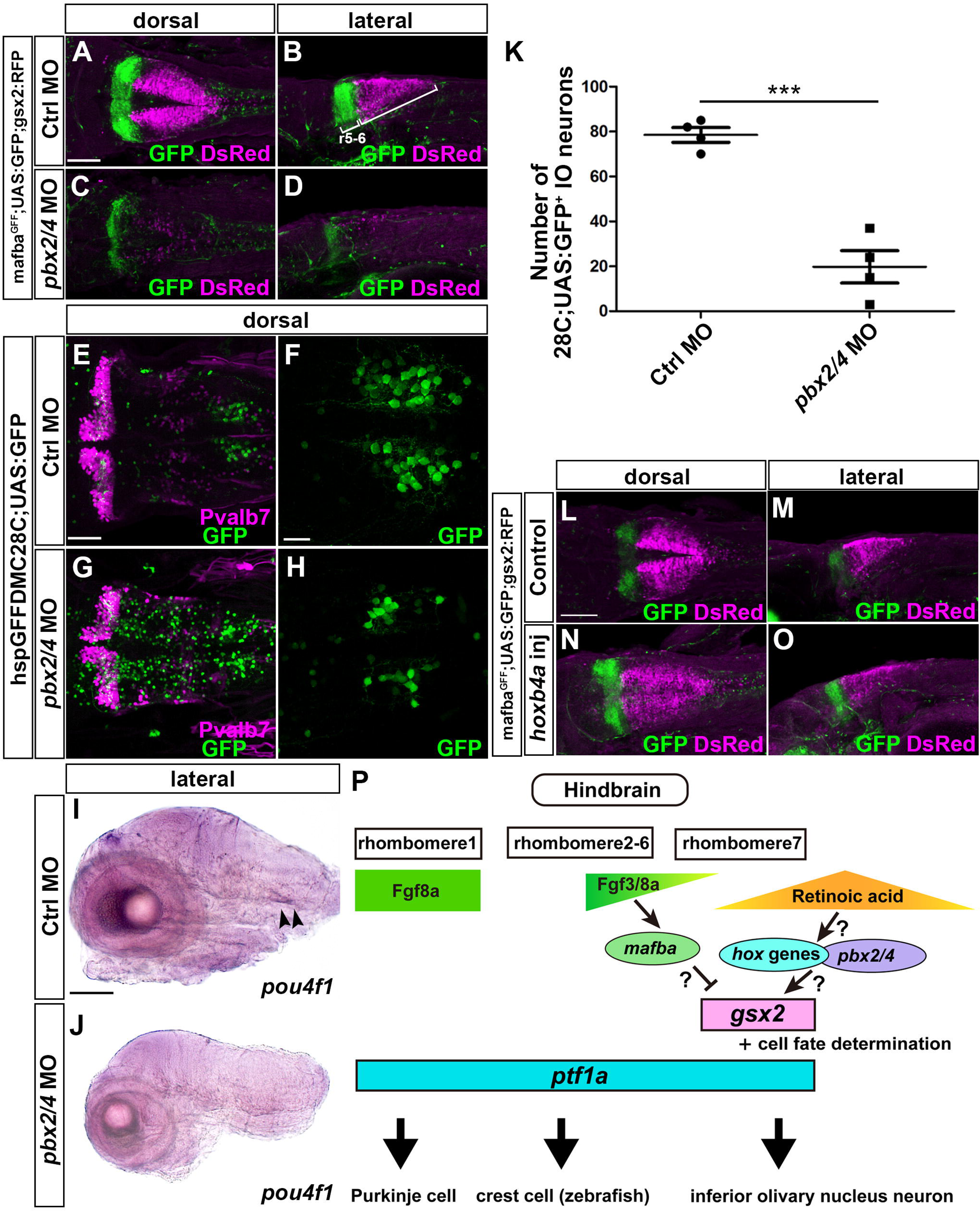
*hox* genes are involved in *gsx2* expression and IO neuronal development. (A-D) 3-dpf control morphant (A, B, *n*=4) and *pbx2/4* morphant (C, D, *n*=4) mafba^GFF^;UAS;GFP;gsx2:RFP larvae were stained with anti-RFP (magenta) and anti-GFP (green) antibodies. (E-H) 5-dpf control morphant (E, F, *n*=4) and *pbx2/4* morphant (G, H, *n*=4) 28C;UAS:GFP larvae stained with anti-Pvalb7 (magenta) and anti-GFP antibodies. (I, J) Expression of *pou4f1* in 5-dpf control morphant (I) and *pbx2/4* morphant (J) larvae. The *pou4f1* signal is marked by arrowheads. (K) Numbers of 28C;UAS:GFP^+^ IO neurons in control and *pbx2/4* morphant larvae. (L-O) 3-dpf control (L, M, *n*=4) and 25 pg *hoxb4a* RNA-injected (N, O, *n*=4) larvae were stained with anti-RFP (magenta) and anti-GFP (green) antibodies. (P) Schematic diagram of the roles of Gsx2 in IO neuronal development. Scale bars: 100 µm in A (applies to A-D), L (applies to L-O) and I (applies to J); 50 µm in E (applies to E and G); 20 μm in F (applies to F and H).

## DISCUSSION

### Expression of *gsx2* and *ptf1a* in IO progenitors located in rhombomere 7

*gsx2* was co-expressed with *ptf1a* in the dorsal ventricular zone of r7 (Fig. 1). Tracing with the Tg lines revealed that *gsx2*- and *ptf1a*-expressing cells (gsx2:RFP^+^ and ptf1a:GFP^+^ cells) gave rise to IO neurons (Fig. 1). The *gsx2* and *ptf1a* mutations abrogated IO neuronal development (Fig. 2, 3). These data indicate that *gsx2/ptf1a*-expressing cells are the neural progenitors of IO neurons. Previous lineage tracing with the Cre-loxP system in mice revealed that IO neurons were derived from *Ptf1a*-expressing neural progenitors (Yamada et al., 2007). Avian grafting studies suggested that IO neural progenitors were located in r6-8 (Ambrosiani et al., 1996; Cambronero and Puelles, 2000; Kawauchi et al., 2006). Our data clearly indicate that in zebrafish, IO neurons are derived from *gsx2/ptf1a*-expressing neural progenitors in r7.

### Role of Ptf1a in development of hindbrain neurons

*ptf1a* (*Ptf1a* in mice) is expressed in neural progenitors in the ventricular zone of the entire dorsal hindbrain in both mice and zebrafish (Fujiyama et al., 2009; Hoshino et al., 2005; Kani et al., 2010; Yamada et al., 2007). Zebrafish *ptf1a* mutants showed a reduction in the number of PCs and crest cells, and the loss of IO neurons (Fig. 3, S4). In mice, *Ptf1a* mutants showed a complete loss of PCs (and other cerebellar GABAergic interneurons), DCN neurons, and IO neurons (Fujiyama et al., 2009; Hoshino et al., 2005; Yamada et al., 2007). Although the mice and zebrafish phenotypes are slightly different, our and their data suggest that *ptf1a*-expressing progenitors give rise to PCs and IO neurons in both species, and also contribute to development of DCN neurons in mice and to crest cells in zebrafish (Fig. 9P).

A milder reduction of PCs in zebrafish suggest that other genes compensate for the lack of Ptf1a. In mice, bHLH-type proneural genes *Neurogenin*(*Neurog*)*1/2* and *Ascl1* are expressed in the ventricular zone of the dorsal hindbrain (Kim et al., 2008; Landsberg et al., 2005; Lundell et al., 2009; Zordan et al., 2008). In mice, *Neurog2* mutants show a reduction in number and defective dendritogenesis of PCs (Florio et al., 2012) while the *Ascl1* mutation led to a reduction of cerebellar GABAergic interneurons and oligodendrocytes (Grimaldi et al., 2009; Sudarov et al., 2011), suggesting the involvement of other proneural genes in the development of PCs and other interneurons in the cerebellum. Zebrafish has *neurog1*, *ascl1a*, and *ascl1b* but not *neurog2* (ZFIN, https://zfin.org). Some of these proneural genes may redundantly function with *ptf1a* in the development of PCs. Unlike PCs and crest cells, IO neurons were completely absent in *ptf1a* mutants (Fig. 3), indicating that Ptf1a is indispensable for IO neuronal development in both mice and zebrafish.

In addition to PCs, crest cells in MONs were reduced in *ptf1a* mutant larvae (Fig. S4). In mice, inhibitory neurons in DCNs are derived from *Ptf1a*-expressing neural progenitors in the r2-5 ventricular zone (Fujiyama et al., 2009). Although zebrafish do not have DCNs, there is a similarity between DCN and MON. Both DCN and MON receive input from hair cells (in the cochlear organ in mice or via the lateral line system in teleosts) (Yamamoto and Ito, 2005). The DCN and MON have a neural-circuit structure similar to the cerebellum and are thus called cerebellum-like structures (Bell et al., 2008; Oertel and Young, 2004). Although crest cells are excitatory, our data suggest that both crest cells and DCN inhibitory neurons are derived from *ptf1a*-expressing progenitors. It is tempting to speculate that the DCN and MON were derived from a common ancestral origin during evolution.

### Role of Gsx2 in the development of IO neurons

The *gsx2* mutants showed a strong reduction or loss of IO neurons (Fig. 2), indicating that Gsx2 is required for IO neuronal development. Previous studies in mice showed that *Gsx2* is expressed in the ventricular zone of the ventral telencephalon, hindbrain and spinal cord, and that it is required for the development of striatal neurons in the telencephalon, olfactory interneurons in the olfactory bulb, and dorsal interneurons in the spinal cord (Corbin et al., 2000; Kriks et al., 2005; Szucsik et al., 1997). Therefore, although Gsx2 is involved in the development of different types of neurons, it plays a similar role in the specification of neurons from their progenitors in the ventricular zone of these brain regions.

In some *gsx2* mutants, a small number of IO neurons remained, and projected CFs to PCs (Fig. 2), suggesting the presence of genes that redundantly function with *gsx2* in IO progenitors. *gsx1*, which is a homologue of *gsx2*, is expressed in the hindbrain (r1-7) and spinal cord of zebrafish (Begemann et al., 2001). In mice, *Gsx1* and *Gsx2* redundantly function in LGE patterning and in specification of dorsal interneurons of the spinal cord (Mizuguchi et al., 2006; Yun et al., 2003). Although the expression domains of *gsx1* and *gsx2* are different in the spinal cord in zebrafish (Satou et al., 2013), it is possible that *gsx1* is co-expressed with *gsx2* and cooperatively functions in the development of a portion of IO neurons. Future studies on *gsx1* expression and *gsx1* mutants will clarify this issue.

### Different roles of Ptf1a and Gsx2 in IO neuronal development

The expression of *ptf1a* and *gsx2* in the caudal hindbrain was independent during early neurogenesis (Fig. 4). Whereas the *ptf1a* mutants showed an increase in apoptosis in the caudal hindbrain, the *gsx2* mutants did not (Fig. 5), suggesting different roles of *ptf1a* and *gsx2* in IO neuronal development. The phenotype of zebrafish *ptf1a* mutants is consistent with that of mouse *Ptf1a* mutants, which also showed increased apoptosis in the IO region (Yamada et al., 2007). These data imply that a major population of IO progenitors that did not differentiate to neurons died by apoptosis in the absence of *ptf1a* in both mice and zebrafish. However, some cells derived from *Ptf1a*-expressing progenitors (*Ptf1a*-lineage cells) become precerebellar neurons that send mossy fibers in *Ptf1a* mutant mice (Yamada et al., 2007), suggesting a role of *Ptf1a* in fate determination of IO neurons in mice. However, in zebrafish, the *ptf1a/gsx2*-lineage cells (ptf1a:GFP^+^ and gsx2:RFP^+^ cells) were present in the dorsal hindbrain and did not migrate ventrally where mossy fiber neurons are supposed to be located in *ptf1a* mutants (Fig. S4, S5). Therefore, even if some cells derived from IO progenitors undergo a change in fate, they are likely a small population in zebrafish.

In contrast to the *ptf1a* mutants, *gsx2*-lineage cells migrated ventrally from the ventricular zone and some of them differentiated to neurons that extended axons to the cerebellum region in *gsx2* mutants (Fig. S5). These neurons, which are potentially the nucleus commissure of Wallenberg, are located above the IOs and send mossy fibers projecting to GCs, as reported for other teleost species (Xue et al., 2004). However, these fibers were also reduced in *gsx2* mutants (Fig. S5), suggesting that development of these neurons partly depends on *gsx2*. We also detected commissure axons of *gsx2*-lineage cells in the caudo-ventral hindbrain of *gsx2* mutants but not in control larvae (Fig. S5). The loss of *gsx2* may lead to a change in fate from IO neurons to commissure neurons. Our findings suggest that *ptf1a* is involved in neuronal differentiation while *gsx2* is involved in the specification of IO neurons. Ptf1a and Gsx2 cooperatively control the development of IO neurons. Ascl1 and Gsx1/2 coordinately control the specification of dorsal glutamatergic sensory neurons in mice (Mizuguchi et al., 2006). Thus, cooperation between proneural gene(s) and *gsx*-family genes is a common mechanism to coordinately control the differentiation and specification of a subset of neurons from their progenitors.

To further address the cooperative role between *ptf1a* and *gsx2* in IO neuronal development, we expressed *gsx2* in *ptf1a*-expressing progenitors by using ptf1a:Gal4-VP16 and UAS:gsx2 Tg lines (Fig. S6). However, ectopic co-expression of *ptf1a* and *gsx2* did not induce ectopic IO neurons in the rostral hindbrain. These data indicate that the co-expression of *ptf1a* and *gsx2* was not sufficient to induce IO neurons. Other factors may function with Ptf1a and Gsx2 in IO neuronal development. One of the candidate cofactors is a bHLH-type transcription factor Olig3, which cooperates with Ptf1a in IO neuronal development in mice (Storm et al., 2009). Future studies will reveal a set of transcription factors that sufficiently initiate the genetic program of IO neuronal development.

### *gsx2* mediates Fgf and RA signals to specify IO neuronal fate

Inhibition of the signal of *fgf3* and *fgf8a*, which are known to be expressed in r4 and are involved in the specification of r5/6 (Maves et al., 2002; Walshe et al., 2002; Wiellette and Sive, 2004), led to reduced *mafba* expression, increased *gsx2* expression, and a reduction in the number of IO neurons (Fig. 6). The *mafba* mutants showed a slight increase in *gsx2* expression and the number of IO neurons (Fig. 7). Fgf signal-mediated *mafba* expression was previously reported for chicken (Marin and Charnay, 2000) and zebrafish (Ghosh et al., 2018). Collectively, these data indicate that the Fgf signal negatively regulates *gsx2* expression and thereby suppress IO neuronal development; Mafba is at least partly involved in this regulation (Fig. 9P). Fgf8a is involved in the formation of the cerebellum (Reifers et al., 1998). In *fgf8a*- or *fgf3/fgf8a*-defective larvae, the number of IO neurons increased but PCs were absent (Fig. 6). Intriguingly, IO neurons project their axons rostrally to the optic tectum region even in the absence of PCs, which are the targets of CFs (Fig. 6), indicating the presence of a PC-independent axon guidance mechanism for CFs. Future studies will clarify this mechanism. As MafB directly activates *hoxb3* expression in mice (Manzanares et al., 1997), the *mafba-hoxb3* cascade may play a role in repressing *gsx2* expression and IO neuronal identify.

During early neurogenesis, *aldh1a2* is expressed in forming somites in mice, chickens, and zebrafish (Begemann et al., 2001; Berggren et al., 1999; Blentic et al., 2003; Grandel et al., 2002; Niederreither et al., 1997). The treatment of mouse embryos with RA increased IO neurons (Yamamoto et al., 2005) but defective RA signals led to a reduced caudal hindbrain in zebrafish (Begemann et al., 2004; Begemann et al., 2001; Grandel et al., 2002). We showed that the inhibition of RA signals resulted in reduced *gsx2* expression and IO neuronal development (Fig. 8). That data indicates that the RA signal positively controls *gsx2* expression and, subsequently, IO neuronal development. *hox* genes function downstream of RA signals (reviewed in (Nolte et al., 2019)). Defects in the RA signal in zebrafish result in a strong reduction or absence of *hoxb4a* in the caudal hindbrain (Begemann et al., 2004; Begemann et al., 2001; Grandel et al., 2002). *hoxb4a* enhancer-driven YFP expression was detected in IO neurons (Ma et al., 2009; Punnamoottil et al., 2008). *Hoxb4* is a direct target of the RA receptor RAR/RXR complex in chickens (Gould et al., 1998). These data suggest that the RA signal controls caudal hindbrain identity through the expression of *hox* genes, including *hoxb4a*, in the caudal hindbrain. Since *hox* paralogues function redundantly, it is difficult to address the roles of individual *hox* genes. We demonstrated that the loss of Pbx2 and Pbx4, which function as cofactors for rostral Hox proteins harboring a YPWM motif (Johnson et al., 1995; Mann and Chan, 1996; Shanmugam et al., 1997), led to a strong reduction of *gsx2* expression and IO neurons (Fig. 9). Furthermore, misexpression of Hoxb4a, which harbors a YPWM motif, elicited the expansion of *gsx2* expression (Fig. 9). Our findings suggest that r7-expressed *hox* genes, such as *hoxb4a*, function directly downstream of the RA signal to control *gsx2* expression and IO neuronal development (Fig. 9). *pbx2/pbx4*-deficient larvae show defects in the function of most rostral Hox proteins and the loss of r2-6 identity (Waskiewicz et al., 2002). While the number of PCs increased, the number of crest cells was normal in *pbx2/pbx4*-deficient larvae (Fig. 9), suggesting that the specification of PCs and crest cells does not rely on Pbx-interacting rostral Hox proteins. The RA-*hox* cascade controls the specification of IO neurons from *ptf1a*-expressing progenitors by regulating *gsx2* expression (Fig. 9P).

*ptf1a*-expressing neural progenitors give rise to three different neuron PCs, crest cells, and IO neurons following positional signals. Gsx2 mediates Fgf and RA positional signals to specify IO neurons. Future studies will reveal genetic programs for the specification of PCs and crest cells. The identification of targets for Gsx2 and Ptf1a will clarify the mechanisms underlying the development and functionality of IO neurons.

## MATERIALS AND METHODS

### Zebrafish strain

The animal work in this study was approved by the Nagoya University Animal Experiment Committee and was conducted in accordance with the Regulations on Animal Experiments at Nagoya University. WT zebrafish with the Oregon AB genetic background were used. For immunohistochemistry and whole mount *in situ* hybridization, larvae were treated with 0.003% 1-phenyl-2-thiourea (PTU) to inhibit the formation of pigmentation. *Tg(UAS:EGFP)* (nkuasgfp1aTg), *Tg(ptf1a:EGFP)*, *TgBAC(mnx2b:GFF)*, *TgBAC(gsx2:LOXP-Tomato-LOXP-GFP)*, and *TgBAC(ptf1a:GAL4-VP16)* were previously reported and characterized (Asakawa et al., 2013; Asakawa et al., 2008; Parsons et al., 2009; Pisharath et al., 2007; Satou et al., 2013). hspGFFDMC28C is a Gal4 trap line that drives UAS-dependent expression in IO neurons (Takeuchi et al., 2015). gSAlzGFFM35A is a *mafba* mutant in which the *mafba* exon is disrupted by the Gal4 gene trap, and is also known as *mafba^nkgsaizgffm35aGt^* (Asakawa and Kawakami, 2018). The *Tg(UAS-hsp70l;gsx2BLRP-P2A-BirA-P2A-mCherry)* line, which harbors a transgene containing *gsx2*, biotin ligase recognition peptide (BLRP), 2A peptide sequences of porcine teschovirus-1 (P2A), biotin ligase (BirA), and mCherry genes, will be described elsewhere. The *mafba^b337^* and *fgf8a^ti282a^* mutants, known previously as *valentino* and *acerebellar*, respectively, were described elsewhere (Moens et al., 1998; Reifers et al., 1998). The allele names of the *gsx2^Δ5^*, *gsx2^Δ8^*, *ptf1a^Δ4^*, and *ptf1a^+11^* mutants established in this study were designated as *gsx2^nub32^*, *gsx2^nub33^*, *ptf1a^nub34^*, and *ptf1a^nub35^*, respectively in ZFIN (https://zfin.org). The zebrafish were maintained at 28°C under a 14-h light and 10-h dark cycle. Embryos and larvae were maintained in embryonic medium (EM) (Westerfield, 2000).

### Establishment of *gsx2* and *ptf1a* mutants by the CRISPR/Cas9 system

The gRNA targets were designed by the web software ZiFit Targeter 4.2 (http://zifit.partners.org/ZiFiT/) (Hwang et al., 2013; Mali et al., 2013). To generate gRNAs, the following oligonucleotides were used: 5′-TAGGCGGAATTCCACTGCTCAA-3′ and 5′-AAACTTGAGCAGTGGAATTCCG-3′ for *gsx2^Δ5^*; 5′-TAGGTCTTCCGGGACGGGCAGA-3′ and 5′-AAACTCTGCCCGTCCCGGAAG-3′ for *gsx2^Δ8^*; 5′-TAGGAAGAGGCGGAGGCGCATG-3 and 5′-AAACCATGCGCCTCCGCCTCTT-3′ *ptf1a^Δ4^*; 5′- TAGGCGTCAAGCTGCCAACGTC-3′ and 5′-AAACGACGTTGGCAGCTTGACG-3′ for *ptf1a^+11^*. gRNA and *Cas9* mRNA syntheses were performed as previously reported (Nimura et al., 2019). A solution containing 25 ng/µL gRNA and 100 ng/µL Cas9 mRNA or 1000 ng/µL Cas9 protein (Toolgen) was injected into one-cell-stage embryos using a pneumatic microinjector (PV830, WPI). The insertion and/or deletion (indel) mutations on the target region were detected by a heteroduplex mobility assay (Ota et al., 2013). The mutations were confirmed by sequencing after subcloning the target regions amplified from the mutant genome into pTAC-2 (BioDynamics Laboratory Inc.).

### Genotyping

To detect indel mutations, the following primers were used: 5′-GTGCGTATCCTCACACATCCACTCT-3′ and 5′-TGCCATCCTCTGGCAGAACG-3′ to detect the *gsx2^Δ5^* mutation; 5′-CATTGGCATGCACTCTCCCG-3′ and 5′-TGAGGATACGCACAGCGGAC-3′ to detect the *gsx2^Δ8^* mutation; 5′-ACCTCAGAGCTGTCCCCTCACAGA-3′ and 5′-GGCAGCTTGACGCAACTGTT-3′ to detect the *ptf1a^Δ4^* mutation; 5′-GGAGGCGCATGAGGTCTGAAGT-3′ and 5′-GCAGTCCCTCGAAAGCATCG-3′ to detect the *ptf1a^+11^* mutation. To identify gSAIzGFFM35A fish, 5′-CGAGGTAGGAGAAGGGCTGT-3′ and 5′-CTGGAGCGTTTGATGGATACAG-3′ primers were used to amplify about 200-bp PCR products from WT but not gSAIzGFFM35A genomic DNA. Since *Tg(gsx2:loxp-DsRed-loxp-GFP)* are BAC Tg fish and contain a portion of the *gsx2* exon1, we could not distinguish the WT allele of *gsx2* from the endogenous locus or the transgene in this fish with the primers described above. To detect WT and mutant alleles of *gsx2* in *gsx2^Δ^*^5^**;*Tg(gsx2: LOXP-Tomato-LOXP-GFP)* fish, the following primers were used: 5′-CCATCAGCATCTCGCTCAG-3′ and 5′-CAGTGAAGCCTTGTCCCTCG-3′. After PCR amplification, PCR products were digested with *Xho*I. The WT *gsx2* allele gave rise to two bands, 21- and 257-bp fragments, while the mutant allele displayed a single band since it cannot be digested with *Xho*I. To detect the *fgf8a^ti282a^* mutation, 5′-CAGGAGGGGGAAACTGATTGTCTAG-3′ and 5′-CCCTTTCTAGGTGGGATTCTTCTC-3′ primers were used. After PCR amplification, PCR products were digested with *Xba*I. The WT PCR products could not be digested whereas the mutant PCR products gave rise to 130- and 20-bp DNA fragments.

### Injection of antisense morpholino oligonucleotides

*fgf3*, *aldh1a2*, *pbx2*, and *pbx4* MOs were translation-blocking antisense MOs that were previously reported (Begemann et al., 2001; Maves et al., 2007; Wiellette and Sive, 2004). One nL of 0.3 mM *fgf3* MO, 0.1 mM *aldh1a2* MO, 0.03 mM *pbx2* MO2, 0.06 mM *pbx2* MO3, 0.06 mM *pbx4* MO1, or 0.06 mM *pbx4* MO2 were injected into one-cell-stage embryos. As a control, we used a standard control MO 5′-CCTCTTACCTCAGTTACAATTTATA-3′. These MOs were obtained from Gene Tool (Philomath, OR, USA).

### Treatment with chemical inhibitors

DEAB (Wako) and SU5402 (Calbiochem) were dissolved in DMSO at 100 mM and 20 mM, respectively. Embryos were treated with 0.25 µM DEAB or 200 µM SU5402 in 0.003% PTU/EM from 6 to 22 h post fertilization (hpf).

### cDNA cloning

Total RNA was isolated from 5- or 12-dpf zebrafish larvae using TRI Reagent (Molecular Research Center, Inc.). cDNA was generated using ReverTra Ace (Toyobo). DNA fragments for *gsx2* or *foxp2* were amplified from the 5-dpf cDNA library using the primers 5′-GACTCTTTGATTATCAAGGATCCCG-3′ (F) and 5′-CGTCTTCTGAGCGCGGATAAT-3′ (R) for *gsx2*, and 5′-CCATGGAGGATAATGGGATG-3 (F) and 5′-TGAGGTAAATTTGGGGGTGA-3′ (R) for *foxp2*, and subcloned to pGEMT-easy (Promega) and pTA2 (Toyobo), respectively (pGEMTe-*gsx2* and pTA2-*foxp2*). Three DNA fragments for *grm5a* (*grm5a*-A, *grm5a*-B, and *grm5a*-C) were amplified from zebrafish genomic DNA with the primers 5′-CACTTTTCTCCGTCCACCAT-3′ (F) and 5′-GCGATCTGGGGAATATTGAA-3′ (R) for *grm5a*-A, 5′-ACCTTCAGTGGGGAGATCCT-3′ (F) and 5′-GATGAAGAGTGCCACGATGA-3′ (R) for *grm5a*-B, and 5′-CGGCCATTATCAAACCATTC-3′ (F) and 5′-TGACGGTAGGATGGTGAACA-3′ (R) for *grm5a*-C, and subcloned into pTAC2 (pTAC2-*grm5a*-A/B/C). Three DNA fragments for *pou4f1* (*pou4f1*-A, *pou4f1*-B, and *pou4f1*-C) were amplified from zebrafish genomic DNA with the primers 5′- GCGATGAGCTGAGATGAGAG-3′ (F) and 5′-AGCTCGAGTGCAGAGTTGTG-3′ (R) for *pou4f1*-A, 5′-GGGTAAGAGTCACCCGTTCA-3′ (F) and 5′-AGTCCGTTGTTGACGAGTCC-3′ (R) for *pou4f1*-B, 5′- CAATTAACGACTCGGACACG-3′ (F) and 5′-TCAGCTATAGCCGCGATTTT-3′ for *pou4f1*-C, and subcloned to pTAC2 (pTAC2-*pou4f1*-A/B/C). The full-length cDNA for *hoxb4a* was amplified from the 12-hpf cDNA library with the following primers: 5′-GGAATTCATGGCCATGAGTTCCTATTT-3′ (F) and 5′-GCTCTAGACTATAGACTTGGCGGAGGTCC-3′ (R). The DNA fragment was digested with *EcoR*I and *Xba*I, and subcloned into pCS2+MT (pCS2+MT-*hoxb4a*).

### *In situ* hybridization

Whole mount *in situ* hybridization was performed as previously reported (Bae et al., 2009). Detection of *ptf1a* and *pou4f1* riboprobes was described previously (Bae et al., 2009; Kani et al., 2010). A digoxigenin (DIG)-labeled riboprobe was made from pTAC2 -*grm5a*-A/B/C, pTA2-*foxp2*, pGEMTe-*gsx2*, and pTAC2-*pou4f1*-A/B/C, using SP6, T3, or T7 RNA polymerase after digestion with restriction enzymes. For *grm5a* and *pou4f1*, a mixture of three riboprobes was used (*grm5a* in Fig. 2, 3; *pou4f1* in Fig. S3). To detect transcripts in the IO neurons of 5-dpf larvae, the head regions rostral to the spinal cord of larvae were cut and used for *in situ* hybridization. To stain larval heads, they were hybridized overnight at 65°C for three nights and incubated overnight with 1/2,000 alkaline-phosphatase conjugated with anti-DIG Fab fragment (Roche) at 4°C for three nights. BM-purple signals were acquired using an AxioPlan-2 microscope equipped with an AxioCam CCD camera (Zeiss).

### Immunohistochemistry

For immunohistochemistry, anti-parvalbumin7 (1/1000, mouse monoclonal, ascites) (Bae et al., 2009), anti-GFP (1/1000, rat, Nacalai Tesque, Cat# 04404-84), anti-DsRed (1/1,000, rabbit, Clontech Laboratories, Inc., Cat# 632496), and anti-cleaved caspase3 (1/500, rabbit, Cell Signaling Technology, Cat# 9661) were used. Alexa Fluor 488 goat anti-rat IgG (H + L, Thermo Fisher Scientific Cat# A-11006), CF488A goat anti-rat IgG (H + L, Biotium Cat# 20023), CF488A goat anti-mouse IgG (H + L, Biotium Cat# 20018-1), CF568 goat anti-mouse IgG (H + L, Biotium Cat# 20301-1), and CF568 goat anti-rabbit IgG (H + L, Biotium Cat# 20103) were used as the secondary antibodies. Larvae were immunostained as described previously (Bae et al., 2009). Some fixed samples were optically cleared with SeeDB reagent as previously reported (Ke et al., 2013; Ke and Imai, 2014). An LSM700 confocal laser-scanning microscope was used to obtain fluorescence images.

### mRNA injection

To make *hoxb4a* mRNA for expression, pCS2+MT-*hoxb4a* was linearized with *Not*I and transcribed with SP6 RNA polymerase in the presence of a G(5′)ppp(5′)G RNA cap structure analog (New England BioLabs Inc.). 25 pg of *hoxb4a* mRNA was injected into one-cell-stage embryos.

### Statistics

Data were analyzed for statistical significance by Student’s *t* test and one-way ANOVA using Graphpad PRISM (ver. 5.1).

## Acknowledgements

We thank Shin-ichi Higashijima and the National Bioresource Project for providing the transgenic zebrafish, Makoto Kobayashi for the *mafba^b337^* mutant, Masato Kinoshita and Feng Zhang for the hSpCas9 plasmids, Kuniyo Kondoh and Yumiko Takayanagi for managing fish mating and care. We also thank the members of the Hibi Laboratory, Hanako Hagio and Naoyuki Yamamoto, for helpful discussion. This work was supported by JSPS KAKENHI Grant Number JP15H04376, JP18H02448 (to M.H.), JP18K06333 (to T.S.), and JP16K18548 (to. M.T.), and CREST Japan Science and Technology Agency (JST) JPMJCR1753 (to M.H.).

## Supplementary material

**Fig. S1. Structure of WT and mutant Gsx2 and Ptf1a.** (A) Structure of WT and mutant Gsx2 and nature of *gsx2* mutations generated by the CRISPR/Cas9 method. The position of the CRISPR/Cas9 target is also shown. Gsx2 has a homeodomain. The *gsx2^Δ5^* mutant contains a 5-bp deletion causing a flame shift that results in 82 unrelated amino acids and a stop codon. The *gsx2^Δ8^* mutant contains an 8-bp deletion causing a flame shift that results in 116 amino acids and a stop codon. The putative mutant proteins lack the homeodomain. (B) Structure of WT and mutant Ptf1a and nature of *ptf1a* mutations generated by the CRISPR/Cas9 method. Ptf1a has a bHLH domain. The *ptf1a^Δ4^* mutant contains a 4-bp deletion causing a frame shift that results in the introduction of 157 unrelated amino acids and a stop codon. The *ptf1a^+11^* mutant contains a 1-bp deletion and 12-bp insertion causing a frame shift that results in the introduction of 162 unrelated amino acids and a stop codon. The putative mutant protein lacks the bHLH domain.

**Fig. S2. Expression of *gsx2* and *ptf1a* is not affected in *gsx2* and *ptf1a* mutants.** (A-D) Expression of *gsx2* in 2-dpf WT (A, C, *n*=2) and *gsx2* mutant (B, D, *n*=3) larvae. (E-H) Expression of *ptf1a* in 2-dpf WT (E, G, *n*=2) and *ptf1a* mutant (F, H, *n*=3) larvae. Scale bars: 100 µm in A (applies to A-B,E-F) and C (applies to C-D, G-H).

**Fig. S3. Defective development of IO neurons in *gsx2***^Δ8^ **and *ptf1a^+11^* mutants.** (A-D) Expression of *pou4f1* in 5-dpf *gsx2^Δ8/Δ8^* (B, *n*=3), and *ptf1a^+11/+11^* (D, *n*=3), and control WT sibling larvae (A, C, *n*=3). (E-H) 5-dpf *gsx2^Δ8/Δ8^* (F, *n*=5), and *ptf1a^+11/+11^* (H, *n*=5), and control WT sibling larvae (E, G, *n*=5) were immunostained with anti-Pvalb7 (magenta) and anti-GFP (green) antibodies. Arrows and arrowhead indicate CFs and IO neurons, respectively. Scale bars: 100 µm in A (applies to A-D); 50 µm in E (applies to E-H).

**Fig. S4. Reduction of PCs and crest cells in *ptf1a* mutant larvae.** (A-L) 5-dpf WT (A-D, I-J, *n*=4) and *ptf1a^Δ4/Δ4^* (E-H, K-L, *n*=6) *Tg(ptf1a:GFP)* larvae stained with anti-Pvalb7 (magenta) and anti-GFP (green) antibodies. (I-L) Highly magnified views of the boxes in A and E. Arrowhead shows crest cells. Arrow shows migrating GFP^+^ cells that are derived from *ptf1a*-expressing progenitors. (M) Number of Pvalb7^+^ Purkinje cells. (N) Number of Pvalb7^+^ crest cells. Scale bars: 50 µm in A (applies to A-C, E-G); 20 µm in D (applies to D, H) and I (applies to I-L). \*\*\**p* < 0.001 (Student’s *t*-test).

**Fig. S5. Lineage tracing of *gsx2*-expressing cells in *gsx2* mutants.** (A-L) 5-dpf WT (A-D, *n*=13), *gsx2^Δ5/Δ5^* (E-H, *n*=8) and *ptf1a^Δ4/Δ4^* (I-L, *n*=3) gsx2:RFP larvae were stained with anti-Pvalb7 (green) and anti-GFP (magenta) antibodies. Arrows and arrowhead indicate CFs and IO neurons, respectively. (C, G, K) Highly magnified views of IO regions. Asterisks indicate axons from gsx2:RFP^+^ cells, which are potentially the nucleus commissure of Wallenberg. Note that the commissure axons of gsx2:RFP^+^ cells were observed in the caudo-ventral hindbrain of *gsx2* mutants but not in the control or *ptf1a* mutants (G). Scale bars: 50 µm in A (applies to A-B, E-F, I-J) and D (applies to D, H, L); 20 µm in C (applies to C, G, K).

**Fig. S6. Ectopic *gsx2* expression alone does not induce IO neurons.**(A, B, D, E) Expression of *gsx2* in 3-dpf *Tg(UAS:gsx2BLRP-P2A-BirA-P2A-mCherry)* (control, A, D) and *TgBAC(ptf1a:Gal4-VP16);Tg(UAS:gsx2BLRP-P2A-BirA-P2A-mCherry)* (B, E) larvae. The *Tg(UAS:gsx2BLRP-P2A-BirA-P2A-mCherry)* line harbors a transgene containing the open reading frame (ORF) of *gsx2*, biotin ligase recognition peptide (BLRP), 2A peptide sequences of porcine teschovirus-1 (P2A), biotin ligase (BirA), and mCherry genes. Expression of Gsx2 and mCherry can be driven by Gal4-VP16. (C) Expression of mCherry was induced where *ptf1a* was expressed in 3-dpf Tg larvae. (F-G) Expression of *pou4f1* in 3-dpf control (F, *n*=3) and *Tg(ptf1a:Gal4-VP16);Tg(UAS:gsx2BLRP-P2A-BirA-P2A-mCherry)* (G, *n*=3) larvae. Although *gsx2* was ectopically expressed in all *ptf1a*-expressing cells, it did not induce ectopic expression of *pou4f1*. IO: inferior olivary nuclei. Scale bars: 100 µm in A (applies to A-G); 50 µm in C.

